# Human Navigation Without and With Vision - the Role of Visual Experience and Visual Regions

**DOI:** 10.1101/480558

**Authors:** Shachar Maidenbaum, Daniel-Robert Chebat, Amir Amedi

## Abstract

Human navigation relies on a wide range of visual retinotopic cortical regions yet the precise role that these regions play in navigation remains unclear. Are these regions mainly sensory input channels or also modality-independent spatial processing centers? Accordingly, will they be recruited for navigation also without vision, such as via audition? Will visual experience, or the lack thereof, affect this recruitment? Sighted, congenitally blind and sighted-blindfolded participants actively navigated virtual mazes during fMRI scanning before and after navigating them in the real world. Participants used the EyeCane visual-to-auditory navigation aid for non-visual navigation.

We found that retinotopic regions, including both dorsal stream regions (e.g. V6) and primary regions (e.g. peripheral V1), were selectively recruited for non-visual navigation only after the participants mastered the EyeCane demonstrating rapid plasticity for non-visual navigation. The hippocampus, considered the navigation network’s core, displayed negative BOLD in all groups.

Our results demonstrate the robustness of the retinotopic nodes modality-independent spatial role in non-visual human navigation to lifelong visual-deprivation, demonstrating that visual input during development is not required for their recruitment. Furthermore, our results with the blindfolded group demonstrate this recruitment’s robustness even to brief blindfolding, but only after brief training, demonstrating rapid task based plasticity. These results generalize the wider framework of task-selectivity rather than input-modality as a brain organization principle to dorsal-stream retinotopic areas and even for the first time to the primary visual cortex.

**Highlights:**

- Both visual and non-visual navigation recruit retinotopic regions
- After training blindfolded subjects selectively recruit V1 & V6 for navigation
- This holds also for participants with no visual experience (congenitally blind)
- The medial temporal lobe showed non-selective Negative BOLD in all groups

**Declaration of interests:** All authors declare that they have no conflicts of interests.

## Main text

Vision is the dominant and most adapted sense which humans use for navigation (Strelow, 1985, Loomis et al., 1993, Ekstrom, 2015, Ekstrom 2018). Accordingly, as we navigate we recruit a wide network of brain regions that are also considered to be visual (Hartley et al., 2003, Vann et al., 2009, Boccia et al., 2014; Ekstrom, 2015, Stensola and Moser, 2016). As such, visual regions, including even the primary visual cortex, are typically included in models of the wider navigation network (Ekstrom et al. et al. 2017, Kong et al. 2017). However, the precise contribution and role they play during navigation are still unclear, especially given the multi-modal nature of navigation. This is especially true as nearly all previous neuroimaging work on human navigation used mainly visual stimuli. This has limited the ability of researchers to disentangle the perceptual role of these regions from the possible spatial processing roles that they may have as well. Thus, it is unclear whether these regions are merely a sensory input port for the visual information, or whether they are also an integral part of the navigation network, performing specialized modality-independent spatial computations for navigation? Furthermore, how dependent on vision is the development of these regions and within what time frame of navigational training with another modality can they be recruited for processing navigation cues arriving from different sensory modalities, like auditory navigation cues?

It is important to remember that human navigation does not rely only on vision - it is a multi-sensory process, combining input from the visual, vestibular and other sensory channels with internal representations. Humans, as any of our readers can self-attest from their experience of moving in the dark or with their eyes closed, are able to navigate without vision, but their navigational performance is changed, generally for the worse (e..g imagine navigating without vision even in a safe environment you know well, such as from your living room to your bathroom, - you may find that you are able to accomplish this task, but with more difficulty than you would accomplishing this task using the visual modality). Behavioral work has suggested that the lack of vision significantly changes navigation and mobility patterns (Millar 1994; Loomis et al., 2001, Souman et al., 2009, Cappagli et al., 2017), but also that offering access to visual information via another modality may lead to higher similarity between blind and sighted individuals in terms of navigation behavior and strategies employed (reviewed in Schinazi et al. 2016). However, in terms of the brain regions underlying neural infrastructure supporting this behavior, the few previous studies exploring these questions were limited by how to present navigation related stimuli non-visually during neuroimaging. Indeed, nearly all previous neuroimaging work on human navigation used mainly visual stimuli. This has limited the ability of researchers to disentangle the perceptual role of these regions from the possible spatial processing roles that they may have as well. Thus, it is unclear whether these regions are merely a sensory input port for the visual information, or whether they are also an integral part of the navigation network, performing specialized modality-independent spatial computations for navigation? Furthermore, how dependent on vision is the development of these regions, and how quickly can they be recruited for processing navigation cues arriving from different sensory modalities, like auditory navigation cues?

Previous studies attempting to explore the neural correlates of navigation without vision and the dependence of the navigation system on visual experience did not use active first person navigation. These previous work resorted to methods such as a passive route recognition task from an external perspective (Kupers et al., 2010), solving finger mazes (Gagnon et al., 2012) and imagined locomotion (Deutschlander et al., 2009; Horner et al., 2016) leading to contradicting results (reviewed in Schinazi et al., 2016), all tasks which are very different from standard navigation. Thus, it remains unclear what role these regions actually play in non-visual active first-person navigation, and whether visual experience is necessary for these regions’ function to develop.

Results from animal models do not help settle these issues or clarify the role of visual experience in shaping these regions, and at best offer only contradicting results One line of research has shown that the rodent grid-cell signal is disrupted in congenital blindness (Chen et al., 2016) and many place fields do not function correctly when visual information is missing (Quirk et al., 1990) or manipulated (Chen et al., 2013). In line with this stance are general reports of a disrupted neural representation of space without vision (Mittelstaedt et al., 1980; King et al. 2009; King et al. 2014), suggesting that vision is critical for the proper calibration of these systems. A contradictory line of research has found that spatial signals were maintained in cases of congenital visual deprivation, such as the spatial firing of place cells in congenitally blind rats (Hill & Best, 1981; Save et al., 1998) and in rodents before eye-opening (Wills et al., 2014; Muessig et al., 2016) or the maintenance of head direction cells (Tan et al., 2015; Bjerkness et al., 2015). Similarly, bats have different maps for information from different senses, including an echolocation based mapping not reliant on vision (Geva-Sagiv et al., 2015). This second group of studies suggests that these types of spatial cells may not require visual input whatsoever to function properly. Furthermore, it remains unclear how these maps would translate from the cellular level in the hippocampus and entorhinal cortex to the system level of the navigation network throughout the brain at large and at visual regions in particular considering their retinotopic organization (though see the recent results from Saleem et al., in press). Finally, even if the results from animal-models were clearly settled, bridging the gap between animal models and human is not trivial for these questions due to the significant differences in the way our senses interact and the way that humans and these models use vision and their other senses to navigate.

Another question underlying the recruitment of visual regions for navigation without vision is these regions’ cellular organization. Low order visual regions have a retinotopic organization (Hubel & Wiesel 1968, Tootell et al. 1998, Pitzalis et al. 2006), which is thought to mirror its function. While retinotopic maps have been shown to robustly form in these regions in cases of congenital visual impairment (e.g. Morland, et al., 2001, Chouinard et al., 2012, Hoffmann & Dumoulin 2015), and some topographic properties have been demonstrated via functional connectivity in the blind (Striem-Amit et al., 2015), the existence of functional topographic biases such as center-periphery in V1 of the congenitally blind is not trivial. furthermore, will regions which have a retinotopic organization for navigation without vision, a case in which such an organization would not seem beneficial, still be recruited?

Thus, in this study we focus on five interrelated questions on the interaction between the spatial and visual retinotopic networks in the human brain by employing for the first time interactive in-scanner non-visual virtual navigation from a first-person perspective in both sighted vision, blindfolded and a group of individuals without visual experience. Specifically, we ask (1) Are visual regions, with emphasis on regions with known retinotopic properties, recruited by visual virtual navigation? (2) Are these same regions recruited when navigating without vision? (3) Can these regions be plastically recruited based on existing properties? (i.e. after only several hours of training and several hours of visual deprivation) (4) What is the role of visual experience in recruiting these regions and in shaping topographic biases (e.g. foveal vs peripheral biases)? (5) Finally, we explore the effect of non-visual navigation on the other key nodes of the navigation network, with emphasis on the hippocampus.

We explore these questions via the unique prism of comparing in-scanner active navigation with and without vision from a first-person perspective between three groups of individuals with varying levels of visual experience (***sighted, sighted-blindfolded*** and ***congenitally blind***) navigating two different Hebb-Williams mazes (Hebb & Williams, 1946). Before and after experiencing the mazes and auditory interface in a real life size Hebb Williams maze, participants navigated two different virtual mazes (henceforth referred to as Maze 1 and maze 2) while undergoing a fMRI neuroimaging (Fig 1). Non-visual navigation was performed via the EyeCane visual-to-auditory sensory substitution device (real-world described in detail in (Maidenbaum et al., 2014a), virtual in (Maidenbaum et al., 2013), see also Methods and Supp Fig 1). Scans were performed before and after a three-day training session in both real and virtual environments (described in detail in Chebat et al., 2015; 2017) to enable us to explore the effect of inherent connections vs. training and to explore the effects of plasticity.

**Figure 1.**
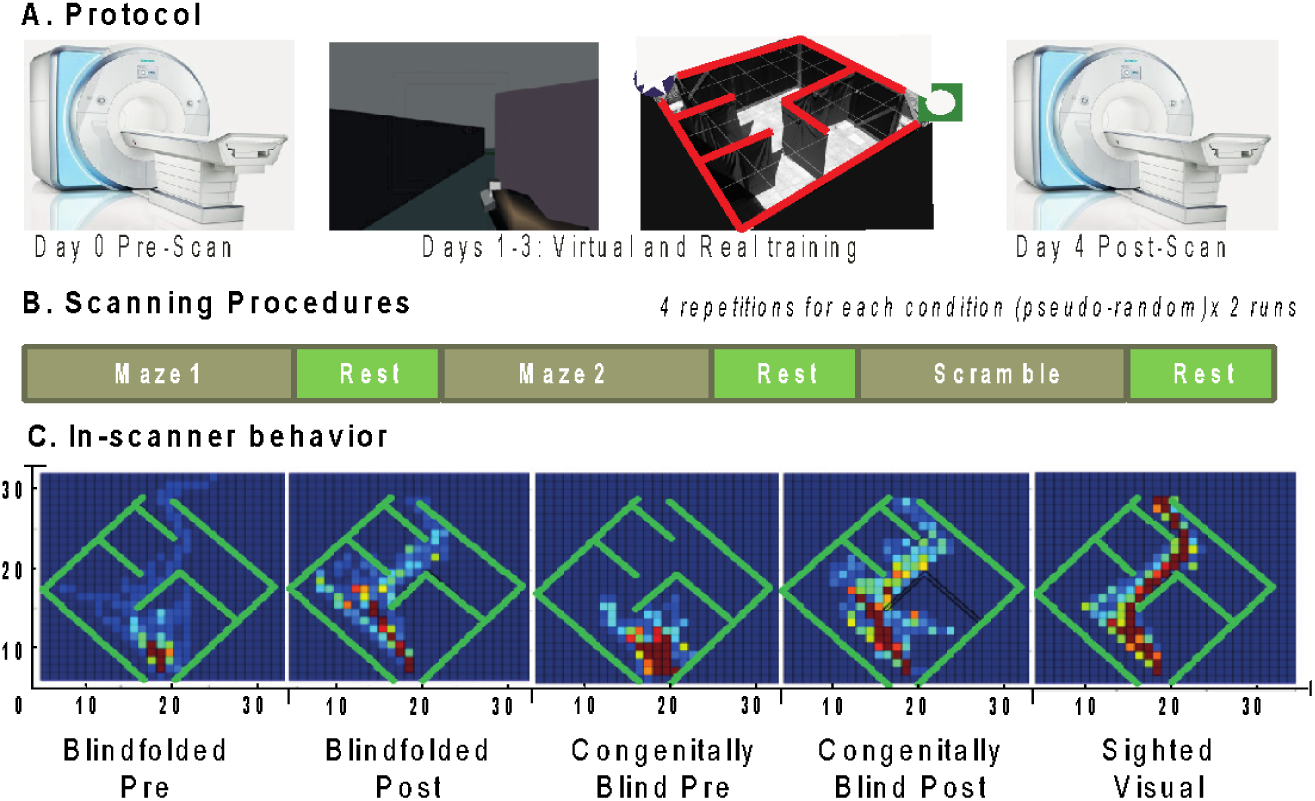
Paradigm and in-scanner behavior. A. *The experimental protocol* consisted of a pre-training fMRI scan, followed by 3 days of training in both real and virtual environments and a post-training scan. B. *The paradigm within the scanner*. There were three types of blocks, Maze 1, Maze 2 and a Scrambled control. Each was repeated 4 times per run, and there were 2 runs on each scanning day. C. *Heatmaps* of the behavior in maze 1 within the scanner for each of the groups. Hotter colors indicate that subjects in that group spent more time in a given location.

## Results

### Subjects can rapidly learn to navigate non-visually

During training outside of the scanner, all participants of the non-visual groups, regardless of prior visual experience and navigation experience, could successfully solve the mazes in both the real and virtual set-ups non-visually with the EyeCane in a way that was not statistically different in terms of errors, collisions and duration from the performance of the sighted who navigated visually (For a detailed description of performance during the longitudinal training see Chebat et al., 2015). Furthermore, after training, non-visual spatial knowledge obtained in one type of environment (real vs. virtual) could be transferred to the other type of environment (from real-to-virtual and virtual-to-real) for all non-visual groups in a matter of hours (Chebat et al., 2017). Within the scanner, in the Pre-training scan subjects actively navigated but could not solve the mazes successfully, while in the Post-training scan they were able to significantly (p<0.003) improve and successfully navigate them (figure 1C). No significant difference was found between the behavioral performance of the two non-visual groups (blind and blindfolded) within the pre-training or within the post-training scans.

### Neuroimaging results

The subjects in the three groups (Sighted, Blind, Blindfolded) were scanned while performing three interleaved conditions - two mazes and a scramble condition. We used two mazes to enable the use of one as a functional localizer for the other in the visual group. The scramble condition included the same audio cues as for when navigating but in a scrambled order and with no connection to the subjects keystrokes, and was used as a control for auditory cues. Each condition was repeated 8 times. To foreshadow the following sections, we found selective recruitment for both visual and non-visual navigation in the dorsal stream (peaking in V6) and in the primary visual cortex (V1).

### Recruitment of dorsal visual regions (peaking at V6/V6A)

We first explored the whole-brain activity in the ***sighted*** group. We performed a whole-brain analysis using the contrast of Maze2>rest and found strong recruitment of the dorsal stream and of the Parieto-Occipital Sulcus, peaking in the Retinotopic Dorsal Precuneus in general, and in particular in visual region V6/V6a. This region is anatomically located at the dorsal end of the parieto-occipital sulcus and has established retinotopic properties (Dechent, et al., 2003, Pitzalis et al. 2006, 2013, 2018).

The functional peak of the dorsal visual stream (thresholded at p<10^-3^) in the previous analysis, corresponding to area V6/V6a, was defined as a functional ROI, and we explored the orthogonal contrast of maze1>scramble to test its stability on the Maze1 blocks. Defining the ROI via an orthogonal contrast is standard procedure for fMRI analysis to avoid double dipping. Accordingly, in the ***sighted*** group maze 2 was used only for the whole-brain analysis and for defining the ROIs, while Maze 1 was used only for the contrast within them.

We found that retinotopic region V6 was recruited for both scramble and navigation in the ***sighted*** group (Betas = 0.2±0.1 and 1.1±0.1 respectively), but with a highly selective preference for navigation (p<10^^-10^) (Fig 2).

**Figure 2.**
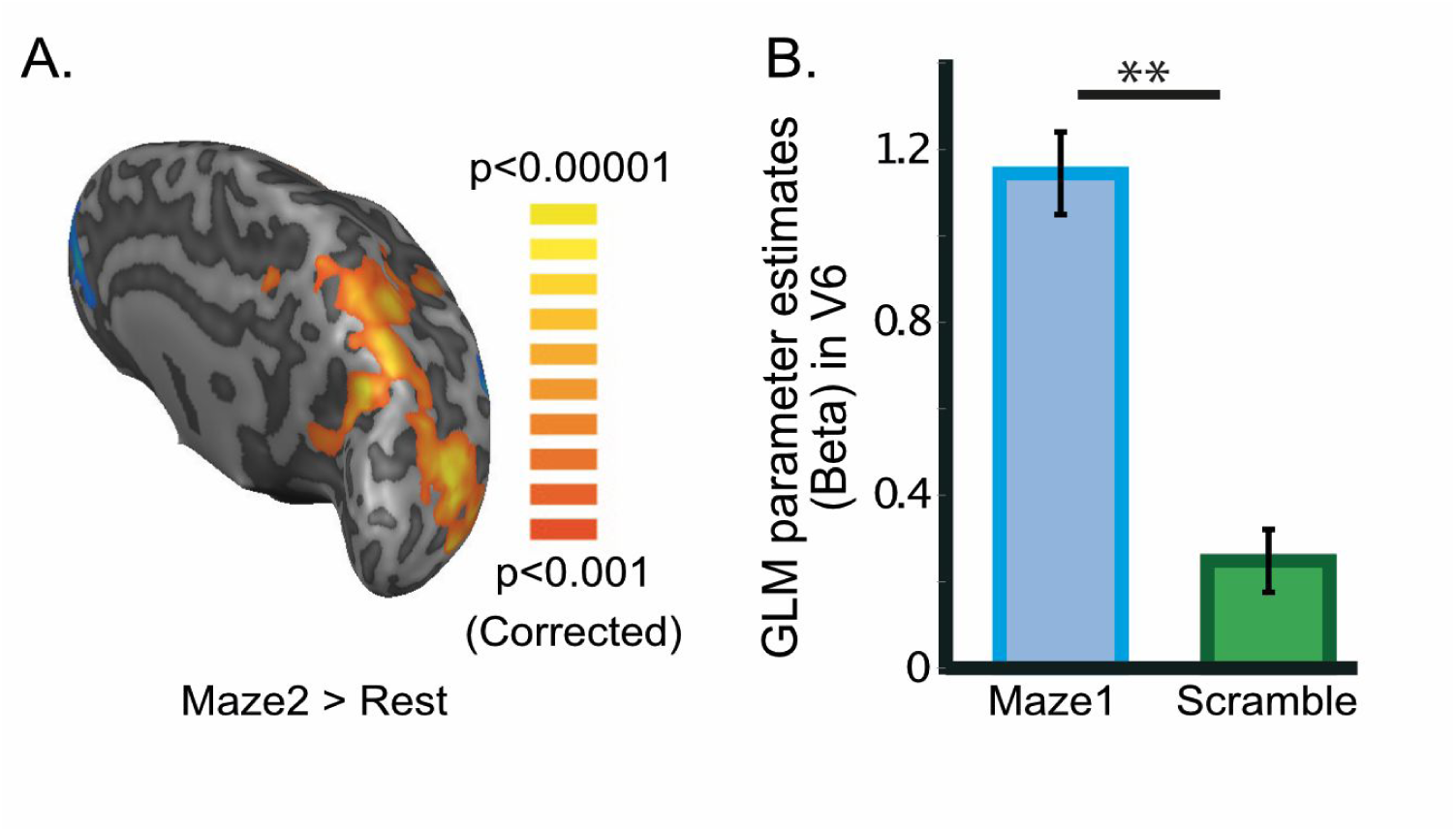
Dorsal stream recruitment in the sighted. Recruitment of the (A) visual dorsal stream for navigation from maze2>rest, (B) Selectivity for navigation peaked in V6\V6a, as demonstrated by comparing maze1 and scramble in the peak ROI found in A. ^**^ denote p<0.01 corrected, Error bars denote the standard error of the mean.

We then tested the ***Blindfolded*** group and found similar results. A whole-brain analysis revealed recruitment of the visual dorsal stream, but not of ventral regions. Using the functional ROI defined above from the sighted group we found that V6\V6a was recruited selectively for navigation both in the pre and post scans (both p<0.01 corrected). However, this selective recruitment for navigation had a 40% increase in the post-training scan, when subjects have learned to use the EyeCane’s auditory input for navigation (Betas = 0.2 and 0.35 respectively, p<0.05 corrected) (Fig 3).

**Figure 3.**
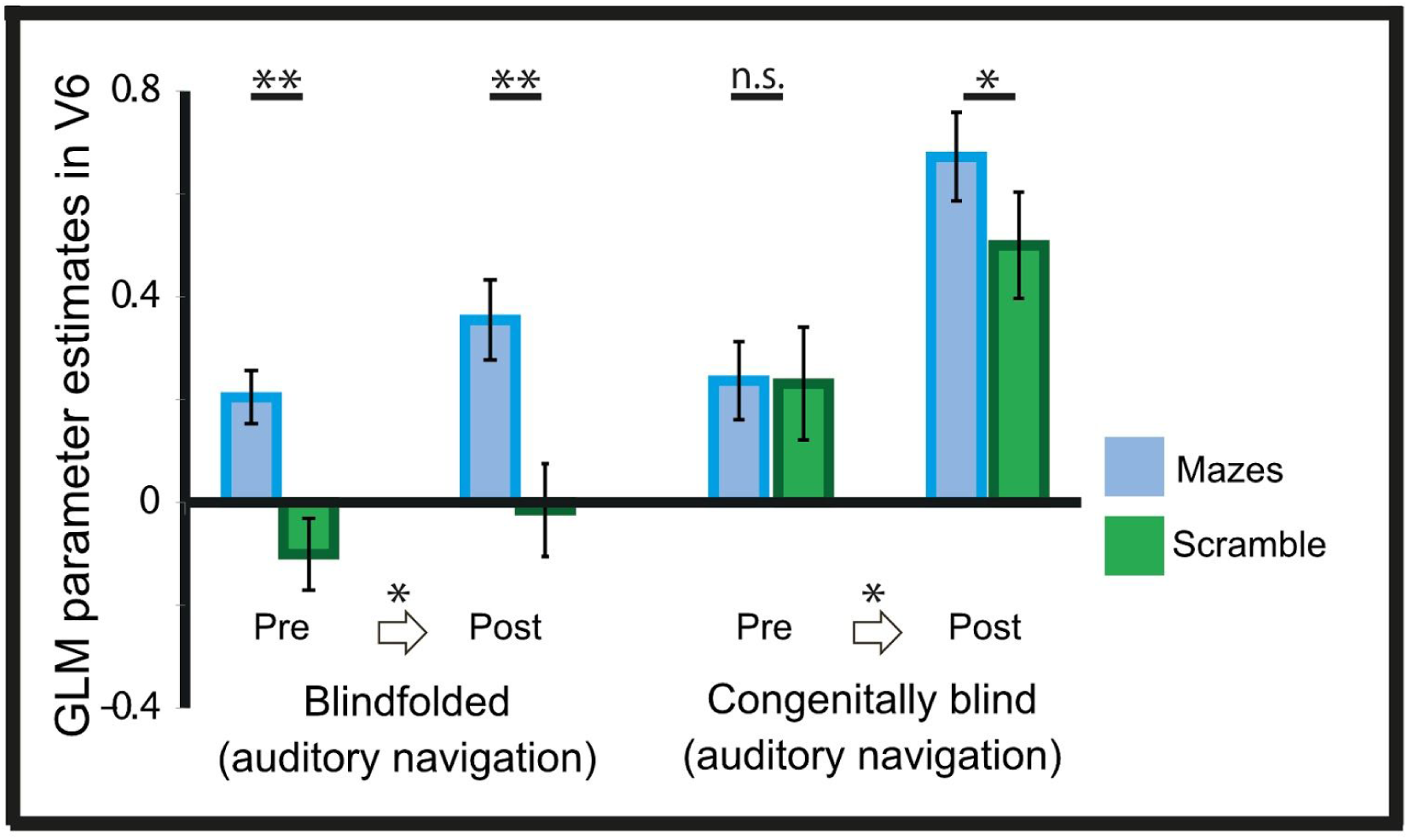
Recruitment of V6 before and after training, demonstrating that post training both groups increase their recruitment and are selective while pre training both groups have low recruitment and the blind group is not selective. ^*^ denotes p<0.05 corrected, ^**^ denote p<0.01 corrected, Error bars denote the standard error of the mean.

This result was robust also in the ***Blind*** group. Whole-brain analysis revealed recruitment of the visual dorsal stream, but not of ventral regions. Using the ROIs defined above from the sighted group for comparison, V6\V6a was recruited un-selectively in the “Pre” (p=0.9), but in the “post” became selective (p<0.05) and its recruitment tripled in size as reflected in the GLM parameter estimate (Betas = 0.23 and 0.67 respectively, p<0.05) (Fig 3).

### Early Sensory Cortices

We then explored the activity in the early sensory cortices. Using ROIs from retinotopy and tonotopy scans derived from an orthogonal group of 15 sighted subjects (taken from Hertz & Amedi, 2010) for A1, foveal V1 and peripheral V1 we found that these areas exhibited different patterns between the pre/post scans and between the groups (Sup Fig 2). ROIs defined by anatomical segmentation of the Calcarine sulcus led to equivalent results.

For the ***sighted*** group, who navigated visually but still heard the same auditory cues as the other groups, A1 showed robust activity (peak of the temporal cortex) but did not show any selectivity (i.e. it was recruited non-selectively for both navigation and scramble) (p=0.7). In contrast to the activity in the auditory cortex, in the primary visual cortex foveal V1 was recruited selectively for scramble (p<10^^-5^) and Peripheral V1 was selectively recruited for navigation (p<10^^-5^) (Fig 4) demonstrating a double dissociation in V1 which fits the classic eccentricity division (Hubel & Wiesel 1968, Tootell et al. 1998).

**Figure 4.**
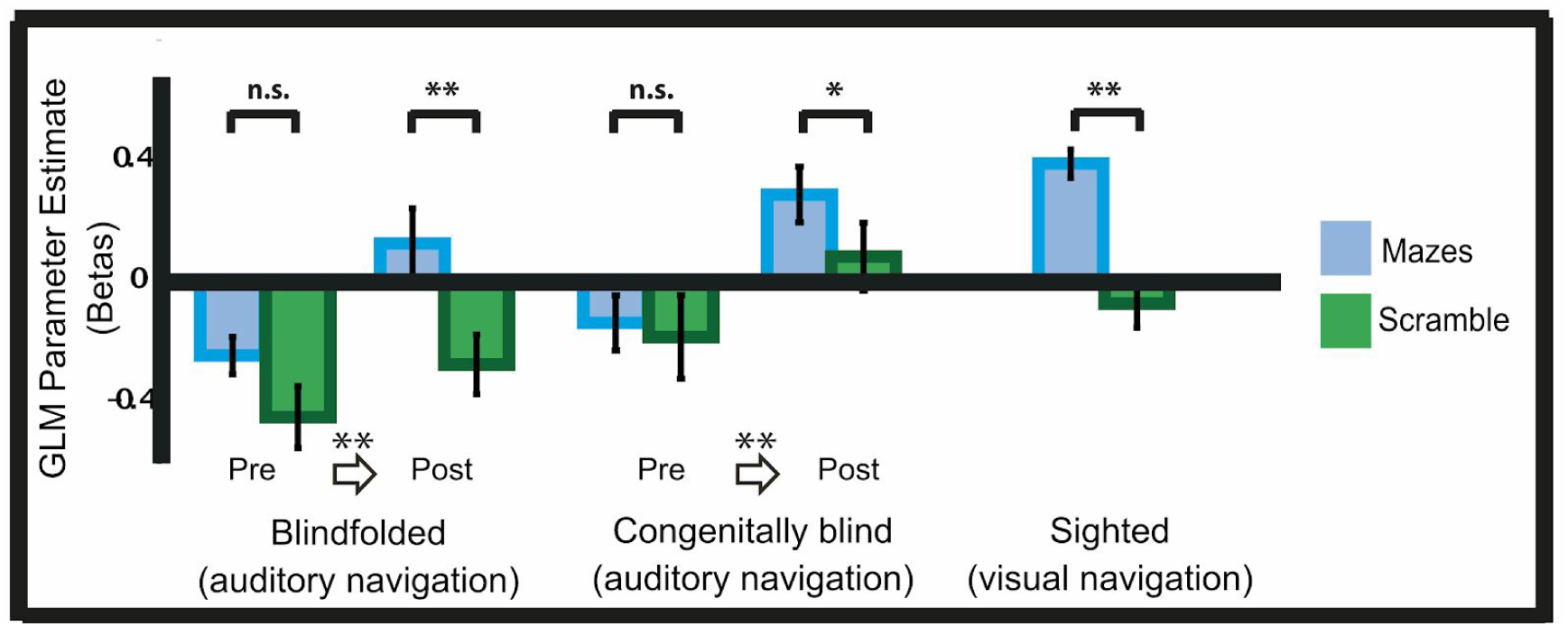
Selective recruitment of peripheral V1 following training with the auditory EyeCane. Before training non-visual navigation does not selectively recruit peripheral V1. Following training peripheral V1 is selectively recruited in both the blindfolded and congenitally blind groups in a pattern similar to the selective recruitment seen in the sighted group.

We then turned to explore these same ROIs in the ***blindfolded*** group. Before training the subjects did not recruit foveal V1, displayed non-selective negative BOLD in peripheral V1 and non-selective recruitment of A1. However, even only after relatively brief training for several hours, both foveal and peripheral V1 were significantly recruited (p<0.01) and both became significantly selective for navigation (p<0.05, p<0.01) (Fig 4). Interestingly, A1 maintained its non-selective recruitment, despite the cues for navigation being auditory!

Finally, we explored these primary sensory cortex ROIs in the ***Blind*** group. Before training subjects recruited foveal V1 and A1 non-selectively and displayed non-selective negative BOLD in peripheral V1. After training however, peripheral V1 was significantly recruited (p<0.01) (Fig 4) and became significantly selective for navigation (p<0.01). Foveal V1 and A1 were not selectively recruited even post training.

### The non-visual nodes of the navigation network

We then returned to the whole-brain analysis to explore activity in the other nodes of the navigation network. We found selective recruitment in several regions including the striatum, with emphasis on the Putamen, and motor regions. On the other hand, we found highly significant negative BOLD in the hippocampus, with emphasis on the posterior hippocampus, for all three groups in both the pre-training and post-training scans (Fig 5).

**Figure 5.**
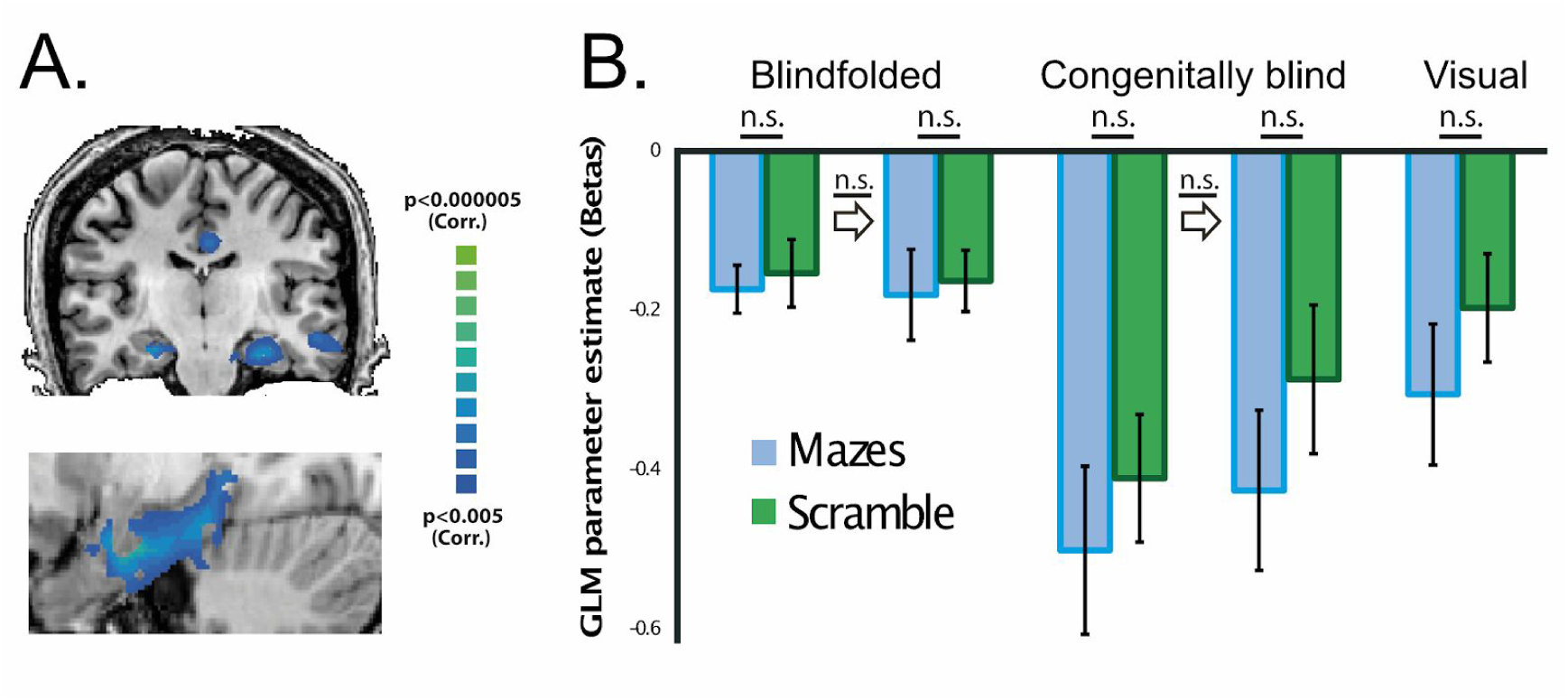
Negative BOLD in MTL regions for all groups, regardless of visual experience or proficiency in the tasks. (A) Whole-brain analysis. (B) Hippocampus ROI.

## Discussion

All groups (***Sighted***, ***Blindfolded***, and ***Blind***) were able to accurately navigate the mazes using the auditory EyeCane interface during the post-training scan, with improved performance vs. the pre-training scan following several hours of training. After learning, all groups, regardless of navigation modality (visual vs. auditory) and visual experience (sighted, blindfolded and congenitally blind), displayed similar general brain activation patterns of wide visual dorsal stream recruitment peaking in V6. Intriguingly all experimental groups/navigation conditions also showed selective recruitment of peripheral V1. Finally, all groups showed negative BOLD in the hippocampus.

Our results demonstrate that (1) on one hand the visual system, stretching from peripheral V1 to dorsal stream areas peaking in V6, includes key nodes recruited for navigation in humans but (2) on the other hand we show that visual experience or visual input is not a critical factor in shaping this selectivity! i.e. the same network can be robustly recruited for navigation without vision and develops independently of visual experience suggesting modality independent task-based computing roles for these regions rather than modality specific ones (Maidenbaum et al., 2014b). The swift recruitment of these regions for a new type of input after only several hours of training, regardless of visual experience or length of deprivation, demonstrates the flexibility and plasticity of these systems. Our results further extend the task machine view of brain organization to dorsal stream retinotopic areas and for the first time also to early topographic retinotopic areas such as the primary visual cortex. This is a key addition to converging evidence from the ventral stream and MT of the blind and high-order temporal cortex of the deaf (reviewed in Amedi et al., 2017).

In the case of the blindfolded group we show very rapid plasticity, occurring within just a few hours of training while blindfolded, can lead to the non-visual recruitment of the visual system. This is surprising as most previous blindfolded perceptual learning studies did not show the recruitment of the visual system in the sighted while using high-resolution sensory substitution devices (Ptito et al., 2005, Pascual-Leone et al., 2005, Kupers et al., 2010), possibly since these complex devices may have required additional training. Additionally, most studies comparing activations of visual cortical areas between sighted and blindfolded groups argued for the necessity of much longer blindfolding durations to recruit visual regions and that even brief exposure to vision would prevent the recruitment of early visual cortex and hamper performance (Pascual-Leone et al., 2005). Here we show for the first time that there is a modulation of the signal following training for sighted blindfolded participants as well as the congenitally blind in both V6/V6A and in the peripheral primary visual areas.

### Task specific sensory independent recruitment of V1

Our results are the first evidence of the selective recruitment of a primary sensory cortex such as V1 for non-visual active navigation in humans. The recruitment was clearly task-based (rather than visual modality based), as the same auditory cues, whether via scramble in the same post-training scan or via navigation blocks in the pre-training task, did not elicit selective recruitment. In stark contrast, A1 did not show any task specific selectivity, despite the fact that all the spatial cues offered to the blindfolded and blind groups were auditory.

From the perspective of the spatial network, these results demonstrate that peripheral V1 is not merely an input node for the brain’s navigation network but rather also performs spatial computations for navigation in and of its own. This result joins recent findings in rodent literature demonstrating spatial based activity in rodent V1 (Saleem et al., in press).

The selective recruitment of peripheral V1, but not of foveal V1, in the congenitally blind group demonstrates that one of the primary visual cortexes’ basic topographic divisions (Levy et al. 2001) can develop in a way that is independent of visual experience. Our results support the view shared with previous research demonstrating that V1 structure may be preserved even in the congenitally blind, who did not have visual input during development (Striem-Amit et al., 2015), and that the right cuneus can be recruited for auditory spatial tasks in the blind (Collignon et al., 2011). This also joins recent work demonstrating the recruitment of foveal, but not peripheral, regions for Braille reading (Siuda-Krzywicka et al. 2016). The emerging picture, matching theories of modality independence (Pascual-Leone et al., 2005, Maidenbaum et al., 2014b, Amedi et al., 2017), is the preservation of the calcarine eccentricity axis for high-resolution tasks in the fovea (e.g. reading faces) and low resolution tasks in the periphery (e.g. spatial perception and navigation).

In the blindfolded group, we cannot rule out the possibility that this recruitment is top-down and visual-imagery based (Muckli et al. 2015), however such an explanation is far less likely for our Blind group who have never had any visual experience.

More broadly, recent years have shown several other examples of the plastic nature of the primary visual cortex in a series of congenital conditions including achiasma, albinism and more (reviewed in Hoffmann & Dumoulin 2015) and in fewer studies even in adulthood (Sabbah et al., 2016) which fit well with our results.

### The dorsal and ventral visual streams

Our results demonstrate a cross-modal recruitment of the visual Dorsal stream, typically characterized as the “information for action” or “how/where” stream (Ungerleider et al. 1994) including its retinotopic regions. This is in line with perceptual tasks exploring both streams directly in the blind (Striem-Amit et al. 2011), or the ventral (Amedi et al., 2001; 2002; Pietrini et al; 2004; Reich et al., 2011; Striem-Amit et al., 2012; Ptito et al., 2012) or dorsal streams (Ptito et al., 2009; Renier et al., 2010; Matteau et al., 2010) independently, and in spatial perception contexts (Kupers et al. 2010).

On the other hand, we find little evidence of ventral stream (“what”, or “perception and recognition”) recruitment in the present study. One might expect the activation to peak in the ventral stream, from spatial areas such as the scene-selective PPA and from there leading to Medial Temporal Lobe (MTL) structures, especially considering the cross-modal recruitment for PPA for perceiving tactile scenes demonstrated by (Wolbers et al. 2011), or in the PHG (Kupers et al., 2010) but we did not find such recruitment. A possible explanation for the lack of ventral stream recruitment may reside in the nature of our task and the level of details of the spatial information – a single pixel representing distance eases navigation notably, but requires much more effort for recognition of landmarks and for other tasks known to recruit the ventral stream.

### Brain organization and function

One of the basic rules of brain organization is that function mirrors organization. This raises an important question – how do we reconcile the spatial representation of our environment offered by spatial cells, such as place, grid and head direction cells, considered required for spatial tasks with the retinotopic organization of visual areas considered to be required for visual processing? Our results show that despite the retinotopic nature of V6 and of the dorsal stream in general, they can be recruited for spatial computations independently of vision (i.e. independent of visual modality or even visual experience). This joins recent results demonstrating non-retinotopic mappings of visual space in the entorhinal cortex (Nau et al., 2018; Julian et al., 2018) highlighting the need for the visual and spatial research communities, who often research the same regions from a different prism and using different terminologies, to better integrate their research.

V6\V6a serves as a good example of this disconnect. On the one hand this region is well established as one of the key nodes of the navigation network, including recruitment for tasks such as first-person perspective visual virtual navigation (Ghaem et al. 1997, Grön et al., 2000, Spiers & Maguire, 2007, Vann et al., 2009, Boccia et al., 2014). This area is typically referred to by spatial researchers as “posterior-parietal”, “dorsal precuneus” and the “parieto-occipital sulcus”. On the other hand, the vision research community defines this area as retinotopic region V6 (Pitzalis et al. 2013), analog to macaque V6 (Galleti et al. 1999) and defines it with clear retinotopic definitions and visual localizers (Pitzalis et al. 2006). This community implicates it in ego-centric visual motion and suggests that the computations it performs support tasks such as subtracting out self-motion signals. These two aspects of V6 fit well together, but each community seems to explore this region separately – what can retinotopic insights offer the spatial view? And how can the spatial tasks be supported by a retinotopic cellular organization?

Our results join the efforts to bridge these gaps by demonstrating that despite the cells in this region being organized by retinotopic principles, this region is sensitive to non-visual egocentric motion and distance calculations, which do not fit trivially into a retinotopic reference frame in a manner independent of visual input.

The interaction between spatial and retinotopic maps is only one example of the wider question of integration across neural representations - e.g. both of these representations need to be translated and integrated also into mappings from our other senses and into motor maps and our brain’s body-centered reference frames (Zaehle et al., 2007, Vann et al., 2009, Gallivan & Culham 2015).

### Why is MTL displaying non-selective negative BOLD?

All groups consistently demonstrated non-selective negative BOLD for navigation in the hippocampus, which is considered the core region for spatial cells and computations in the brain. We initially expected a difference between visual and non-visual navigation in this region, however the results from the visual group matched those from the non-visual groups. Why is this signal negative and non-selective for navigation?

Negative BOLD has previously been reported in these regions for spatial tasks (Shipman and Astur, 2008, Viard et al., 2011), sparking a debate around them and offering several potential directions (Nilsson et al., 2013, Reas and Brewer, 2013a), none of which are wholly satisfactory as a general explanation.

Here, the signals consistency across groups suggests against explanations based on sensory perception (Azulay et al., 2009) or on differences between the groups. Explanations based on task difficulty and levels of task familiarity (Reas and Brewer, 2013b, Hirshhorn et al., 2012) are challenged here by the signal consistency both in the pre and post scans. Explanations based on differential hippocampal blood flow properties changing the BOLD reaction, interaction with neural oscillations or rest-based contrast issues (Stark et al. 2001, Schridde et al., 2008, Nilsson et al., 2013, Ekstrom, 2010; Fellner et al. 2016) are challenged by the existence of many previous tasks in the literature (Grön et al., 2000, Hartley et al., 2003, Hirshhorn et al., 2012), including in the blind (Gagnon et al., 2012), where the hippocampus does exhibit a positive BOLD reaction. Thus, we see task related explanations as the most likely, specifically, egocentric vs. allocentric or spatial-memory vs. response navigation strategies (Iaria et al. 2003). Our task was extremely egocentric as the subject’s main information was the relative distance of a single point compared to their own location. Our task was also possibly reliant on response strategies, as subjects were constantly reacting to the EyeCane distance-information as they searched for openings. This is strengthened by the existence of a strong selective recruitment of the striatum and of the parietal regions in all groups (visual and non-visual, regardless of visual experience) which are considered to support such calculations. However, the existence of the negative signal in the visual groups, where these strategies were less exclusive weakens these directions somewhat as well.

Regardless of the preferred explanation, this negative signal highlights the need for further comparative exploration of the role of other brain regions in navigation beyond MTL in both humans and animal models.

### The use of virtual environments for studying navigation

The immense potential of virtual reality paradigms for neuroscience research still remains largely untapped (Bohil et al. 2011). This powerful tool must be used with some caution however since they are only simulations of real-world navigation, lacking features such as idiothetic and proprioceptive cues.

Recent work in rodents has led to debate concerning the generalizability of results between virtual and real environments (Aghajan et al., 2015, but see Aronov & Tank, 2014). Similarly, several differences between real and virtual environments have been found in humans (Aghajan et al., 2017). On the other hand, there is an extensive literature demonstrating the similarities between virtual and real navigation demonstrating the similarity of the two and the ability to utilize current VR paradigms to understand real navigation (Ekstrom et al., 2018). Thus, while current results from virtual environments are valuable, when future better methods for real-world neuroimaging are available this work should be explored also during real-world navigation.

Here, our subjects only navigated virtually which is a clear limitation on our results. We attempted to address this confound by including real-world versions of the task during training demonstrating similar behavioral patterns (Chebat et al. 2015), and by exploring the behavioral transfer of spatial information and strategy between the virtual and real-world conditions (Chebat et al. 2017).

### General Conclusions

The navigation network, including early retinotopic visual nodes, demonstrates task-specificity, and sensory-independent organization and rapid plasticity. We have shown the potential for fast cross-modal plasticity following brief or lifelong visual deprivation with just several hours of training in a task where blindfolded and blind participants navigated non-visually during neuroimaging. These results shed new light on the navigation network itself and on the role of the visual nodes within it. Furthermore, they demonstrate the potential for cross-modal plasticity and extend the cross-modal task-specific sensory-independent theory of brain organization to the dorsal stream and to low-level topographic sensory cortices.

## Acknowledgements

The Authors wish to thank Maxim Dupli, Natalie Padon, and Maymana Moati for their help in conducting the experiment. Special thanks to the Optical Center of Jerusalem and Mr. Laurent Levy for space for building the mazes and conducting the experiment. This work was supported by a European Research Council grant (grant number 310809); The Charitable Gatsby Foundation; The James S. McDonnell Foundation Scholar Award (grant number 220020284); The Israel Science Foundation (grant number ISF 1684/08); The Edmond and Lily Safra Center for Brain Sciences (ELSC) Vision center. DRC was funded by the Azrieli International Post-Doctoral Fellows program and the ELSC post-doctoral scholarship. SM was funded by a Haubenstock Stiftung award. The funders had no role in study design, data collection and analysis, decision to publish, or preparation of the manuscript.

## Materials and Methods

### Participants

32 subjects participated in this study. Of them, 22 were Sighted; Nine of whom were blindfolded (range: 21-51 years, average: 28 years, mode: 24 years) and Thirteen of whom performed the task using vision (8 women, range: 19-55 years, average: 28 years, mode: 21 years); Ten congenitally blind participants (4 women, range: 23-59 years, average: 36 years, mode: 23 years) all with documented complete blindness from birth (see table 1) completed the experiment, with 2 additional blind participants dropping out during the experiment by their own choice without completing a Post scan. Blindness was peripheral in all cases (i.e. not due to a brain injury). The sighted-visual group was briefly trained on the task prior to entering the magnet and only performed a single scan, while the other two groups underwent three training sessions between scans, spread out over a three day period. This training is described in (Chebat et al., 2015). Demographics of the blind participants are summarized in Table 1. All CB participants were adept white cane users and had previously received training in orientation and mobility. All participants signed informed consent forms. This experiment was approved by the Hebrew University’s ethics committee and conducted in accordance with the 1964 Helsinki Declaration.

**Table 1:** Demographics of the congenitally blind participants

| Name | Age | Sex | Group | Onset | Light Perception | Cause of Blindness |
| --- | --- | --- | --- | --- | --- | --- |
| D.A. | 59 | M | Congenitally Blind | birth | None | Retinopathy of Prematurity |
| M.D. | 23 | M | Congenitally Blind | birth | None | Congenital Glaucoma |
| U.M. | 41 | M | Congenitally Blind | 6-7 months | None | Retinopathy of Prematurity |
| O.B | 38 | M | Congenitally Blind | birth | None | Retinopathy of Prematurity |
| S.S. | 54 | M | Congenitally Blind | birth | None | Retinopathy of Prematurity |
| O.G. | 38 | F | Congenitally Blind | birth | None | Anophthalmia |
| E.D. | 33 | M | Congenitally Blind | birth | None | Retinopathy of Prematurity |
| E.H. | 30 | F | Congenitally Blind | birth | Faint | Retinopathy of Prematurity |
| E.N. | 30 | F | Congenitally Blind | birth | None | Anophthalmia |
| M.S. | 37 | F | Congenitally Blind | birth | None | Anophthalmia |

## The EyeCane sensory substitution device for auditory navigation

The EyeCane (Supp Figure 1) is an in-house minimalistic sensory substitution device, translating distance into auditory and tactile cues in real-time (Maidenbaum 2014a). It has already been used for several other tasks such as distance estimation and obstacle detection & avoidance and we have demonstrated that new users can master its use within 5 minutes of training (Maidenbaum 2014a). We created a virtual version of the EyeCane (Maidenbaum et al., 2013), translating the distance to a single point to auditory cues identical to those presented in the real-world, and used it for a series of tasks, including comparatively exploring mobility patterns and efficiency (Maidenbaum et al., 2014c).

The virtual version of the EyeCane uses a ray-casting algorithm within the virtual environment to calculate if there is a wall in front of the user, and if so at what distance. It then plays the corresponding auditory cue to the one produced by the EyeCan ein the real-world. This cue conveys the distance to the virtual wall via frequency such that the closer the wall the higher the frequency. Spatial information is perceived by sweeping the device to scan the environment, much like a typical white cane is used. This sweeping motion enables the construction of a mental representation of the users’ surroundings.

### The virtual navigation setup

Our mazes were based on the classic Hebb-Williams mazes (Hebb & Williams, 1946), which have been previously used for testing spatial perceptual learning in a wide variety of species, from mice (Meunier et al., 1986) to non-human primates (Zimmermann, 1969), and even in a virtual rendition for humans (Shore et al., 2001). Our virtual environments (Maidenbaum et al., 2013) were created with Blender 2.49, and Python 2.6.2. The location and orientation of the user’s avatar and Virtual EyeCane was tracked at all times at a rate of 60 Hz (identical to the function rate of the virtual environment, thus covering any possible in-game activity) to enable recreation and analysis of the participants’ behavior and were aligned via logged Triggers to the neural data. The environments have a graphical output to the screen, which was used by the ***Sighted*** group. The participants always experienced the environments in the 1st person, and not as a map overview. The virtual mazes matched perfectly the real-world mazes used in training in terms of spatial layout. Distances within the environment were set so that each “virtual meter” correlates to a real-world meter (i.e. the real-world maze was 4.5m, so the virtual maze was set to 4.5 virtual meters, with the same holding for the scale of avatar size and motion).

### Paradigm

In the scanner, subjects underwent 8 repetitions of three conditions:

1. Maze 1 – A virtual recreation of Hebb-Williams maze 3, on which subjects were trained between the pre and post scans
2. Maze 2 – A virtual recreation of Hebb-Williams Maze 6, on which subjects were not trained
3. Scramble – Controlled for auditory stimuli and motor key presses without actual navigation via scrambled visual and auditory stimuli from the mazes, with no effect to subject’s keystrokes (though subjects were told to move normally and were not informed of this).

### fMRI parameters

The BOLD fMRI measurements were acquired in a whole-body 3T GE Signa scanner (GE Medical Systems, USA). We used the standard gradient-echo EPI pulse sequence. We acquired 27 slices of 4.5 mm thickness and 0 mm gap. The data in-plane matrix size was 64X64, field of view (FOV) 220 mm 220 mm, time to repetition (TR) 1.5s, flip angle ¼ 70 and TE 35 ms. The first 10 images of each scan were excluded from the analysis because of non-steady-state magnetization. The fMRI image processing and statistical analysis were performed with the Brain Voyager QX 2.8 software package (Brain innovation; Goebel et al., 2006) using standard preprocessing procedures (identical to Striem-Amit et al. 2012).

### Statistics

All P values were corrected for multiple comparisons via FDR with q<0.05 (Benjamini and Hochberg, 1995). Unless otherwise specified all tests comparing pre-post are two tailed paired t-tests corrected for multiple comparisons.

### Defining the ROIs

Retinotopic area V6 was characterized anatomically, based on its anchoring to the dorsal part of the parieto-occipital sulcus (Pitzalis et al., 2006). The functional ROI used for the analysis in the orthogonal conditions and in the other groups was defined *functionally* as the peak of dorsal-stream activation, which fell within the anatomically defined V6.

Primary sensory areas were defined based on localizers from an external group of 15 subjects. These ROIs have been used before in (Hertz & Amedi 2010). Additional ROIs for fovea and periphery were created based on single-subject level anatomical segmentation of the calcarine sulcus to 3 parts, and definition of the posterior as foveal and the anterior as peripheral. We repeated our analysis for these ROIs and the results were equivalent.

### Data availability

All data is available upon reasonable request from the lead author.

## Supplementary figures

**Sup figure 1.**
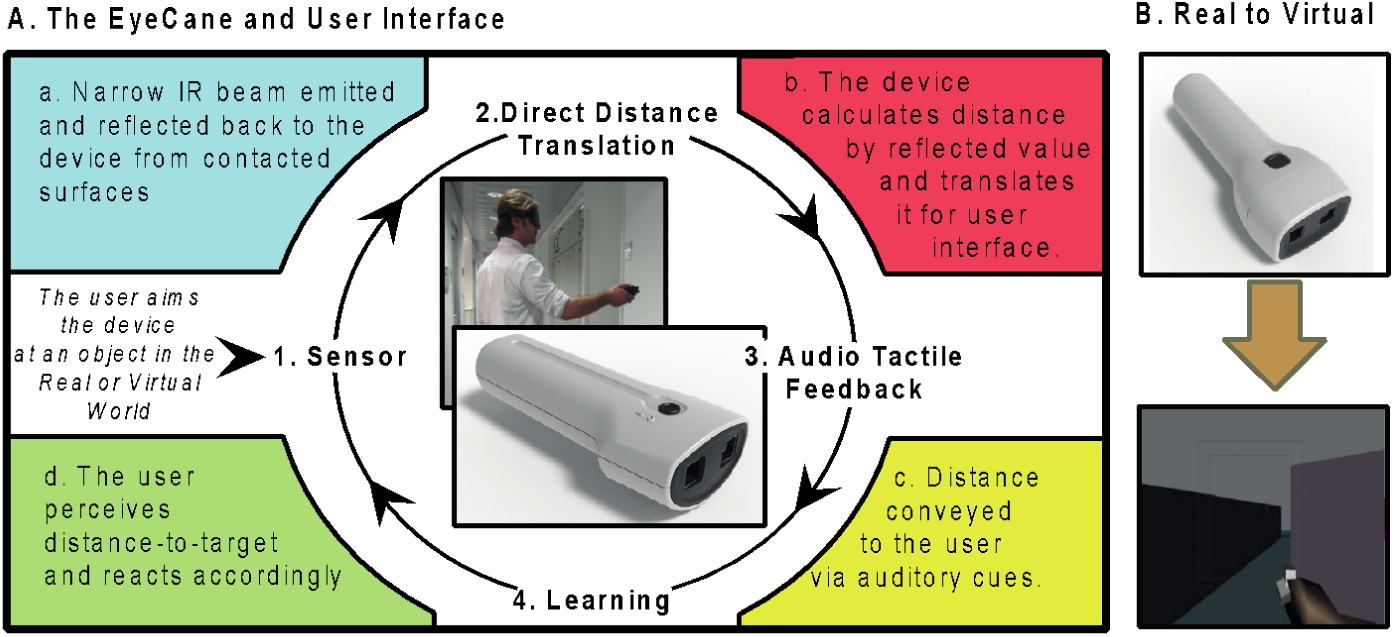
The EyeCane. (A) How to use the EyeCane. (B) Real and virtual versions.

**Sup figure 2.**
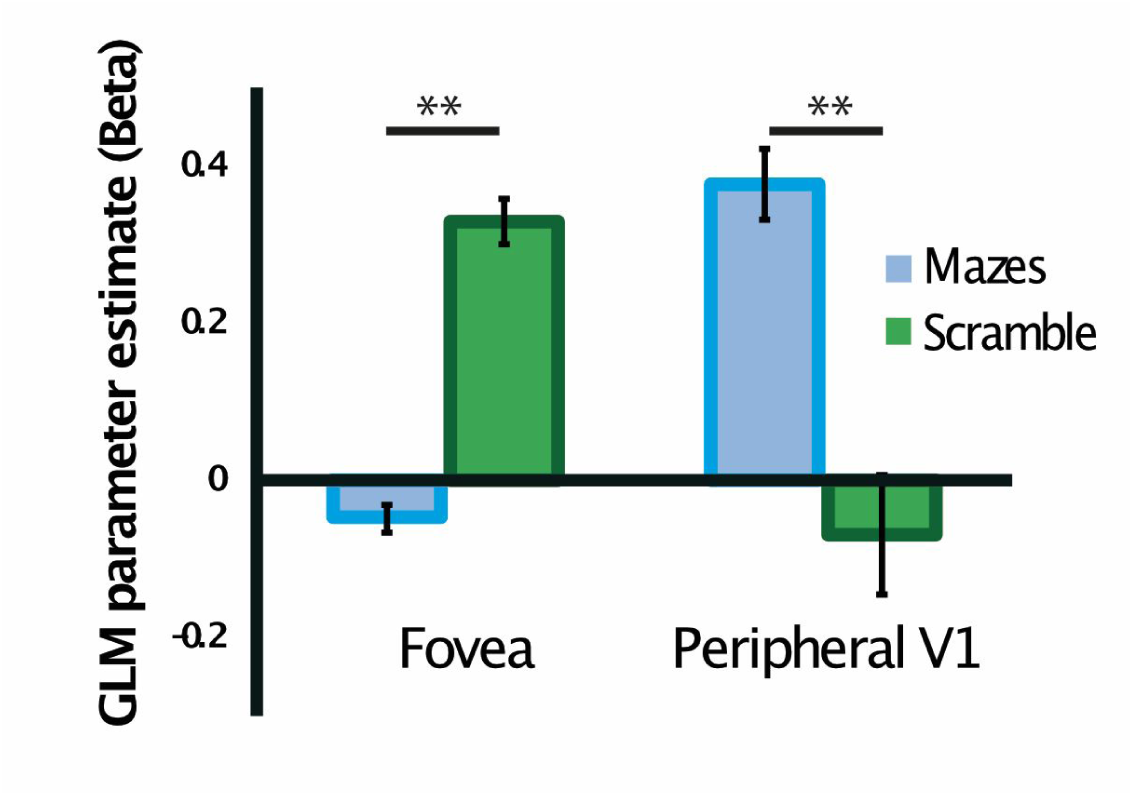
Fovea-Periphery dissociation in the sighted. Double dissociation between Navigation and Scramble in foveal and peripheral V1 of the sighted group navigating visually. ^**^ denote p<0.01 corrected, Error bars denote the standard error of the mean.

